# Snow Buntings, an arctic cold-specialist passerine, risk overheating under intense activity even at low air temperatures

**DOI:** 10.1101/2023.09.11.557251

**Authors:** Ryan S. O’Connor, Oliver P. Love, Lyette Regimbald, Alexander R. Gerson, Kyle H. Elliott, Anna L. Hargreaves, François Vézina

## Abstract

Birds maintain some of the highest body temperatures (T_b_) among endothermic animals. Often deemed a selective advantage for heat tolerance, high T_b_ also limits the capacity to increase T_b_ before reaching lethal levels. Recent thermal modelling suggests that sustained effort in Arctic birds might be restricted at mild air temperatures (T_a_) during energetically demanding life history stages, which may force reductions in activity to avoid overheating, with expected negative impacts on reproductive performance. Consequently, understanding how Arctic birds will cope with increasing T_a_ has become an urgent concern. We examined within-individual changes in T_b_ in response to an experimental increase in activity in outdoor captive Arctic cold-specialised snow buntings (*Plectrophenax nivalis*), exposed to naturally varying T_a_ from -15 to 36 °C. Calm buntings exhibited a modal T_b_ range from 39.9 – 42.6 °C. However, we detected a dramatic increase in T_b_ within minutes of shifting birds to active flight, with strong evidence for a positive effect of T_a_ on T_b_ (slope = 0.04 °C/°C). Importantly, by T_a_ of 9 °C, flying buntings were already generating T_b_ ≥ 45°C, approaching the upper thermal limits of organismal performance (i.e., T_b_ = 45 - 47 °C). Under scenarios of elevated T_b_, buntings must increase rates of evaporative water loss and/or reduce activity to avoid overheating. With known limited evaporative heat dissipation capacities, we argue buntings operating at peak energy levels will increasingly rely on behavioral thermoregulatory strategies (i.e., reducing activity) to regulate T_b_, at the potential detriment to nestling growth and survival.

## Introduction

Birds maintain high and relatively constant body temperatures (e.g., 38-41°C) across a wide range of environmental temperatures (Clarke and Rothery, 2008; Mcnab, 1966; Prinzinger et al., 1991). When heat stressed, birds often allow body temperature to increase above normothermic levels (i.e., facultative hyperthermia), thereby aiding in water conservation by simultaneously augmenting the thermal gradient between body and air temperature and delaying the onset of evaporative water loss (Gerson et al., 2019; Tieleman and Williams, 1999; Weathers, 1981).

However, because birds already regulate high body temperatures, their capacity for increasing body temperature above normothermic levels is limited before reaching potentially lethal body temperatures of 45 – 47°C (Freeman et al., 2020; McKechnie and Wolf, 2019). Consequently, birds working at peak metabolic levels (e.g., nestling-provisioning adults) under high heat loads regularly increase their evaporative heat dissipation behaviours and/or reduce activity to limit endogenous heat production and avoid lethal body temperatures (Clark, 1987; Smit et al., 2016). Individuals are therefore presented with a trade-off during periods of elevated ambient temperatures (T_a_) wherein they must increase thermoregulatory behaviours while decreasing other essential activities, culminating in possible costs on body condition, survival and reproduction (Cunningham et al., 2021; du Plessis et al., 2012). For example, van de Ven et al., (2020) showed that the probability of successful fledging among southern yellow-billed hornbills (*Tockus leucomelas*) fell below 50% when maximum air temperatures rose above 35.1°C. Moreover, nestling hornbills exposed to the hottest weather fledged the nest 50% lighter than those raised under the coolest nesting conditions. Thus, the complex interaction among environmental temperature, activity, and body temperature regulation can have pronounced impacts on the quality of parental care among avian species (see review by (Cunningham et al., 2021).

The Arctic is currently experiencing the fastest rates of warming on Earth (Rantanen et al., 2022). Such rapid change has already affected critical ecological processes (Gilg et al., 2012; Legagneux et al., 2014; Serreze and Barry, 2011), with recorded impacts stemming from phenological mismatches (Carey, 2009; Gaston et al., 2009), increased parasitism (Gaston and Elliott, 2013; Gaston et al., 2002) and invasive species (Gaston and Woo, 2008; Hamer, 2010). In contrast, the direct impacts of rapidly increasing temperatures on the behavior and physiology of Arctic species is lesser known, despite these species being potentially significantly at risk given their evolved traits for conserving heat and coping with extreme cold (Blix, 2016). As such, Arctic cold-speclialists may now be facing signifcant performance constraints in a rapidly warming Arctic due to co-adaptations to perform optimally under cooler conditions, but at the expense of reduced performance at warmer temperatures (Angilletta and Cooper, 2010; Boyles et al., 2011b). Indeed, recent research among Arctic birds suggests comparatively high heat sensitivities, with individuals exhibiting signs of heat dissipation at relatively moderate air temperatures and/or low body temperatures (Choy et al., 2021; O’Connor et al., 2021).

Moreover, the capacity for physiological thermoregulation through increases in evaporative water loss among Arctic birds appears significantly constrained (Choy et al., 2021; O’Connor et al., 2021). Indeed, recent thermal modelling work has predicted that under increasing air temperatures, cold-adapted Arctic birds will have to allocate more time toward regulating body temperature through behavioral adjustments to avoid lethal hyperthermia, but at the expense of interfering with important life-history stages (e.g., reproduction) (O’Connor et al., 2022).

Snow buntings (*Plectrophenax nivalis*) are an Arctic songbird that spends most of their life in sub-zero air temperatures (Le Pogam et al., 2021a; Meltofte, 1983). Even on their breeding grounds, buntings can experience blizzard conditions and air temperatures of -25°C (Meltofte, 1983). Consequently, buntings are known cold-specialists, shown to tolerate air temperatures equivalent to -90 °C under laboratory conditions (Le Pogam et al., 2020). In contrast, O’Connor et al., (2021) recently showed that snow buntings are considerably constrained in their capacity to dissipate heat through evaporative pathways and become heat stressed at relatively moderate air temperatures. Consequently, under increasing heat loads buntings must rely on behavioural thermoregulatory strategies to maintin non-lethal body temperatures (O’Connor et al., 2022). To investigate body temperature patterns and heat dissipation behaviours in buntings, we worked with an outdoor captive population of snow buntings and took advantage of the natural seasonal variation in air temperatures from -15 °C (winter) to 36 °C (summer). We measured body temperature patterns in calm birds to establish baseline inter-individual variation in body temperature, which allowed for determining individual spare capacity for hyperthermia when active. We also induced flight activity to quantify increases in body temperature relative to air temperature and observe any resulting heat dissipation behavior (e.g., panting or wing spreading). We expected maximum body temperatures in actively flying buntings to increase with air temperature, with buntings showing signs of heat stress via panting at comparatively low air temperatures. We were particularly interested in determining at which T_a_ birds would begin to increase T_b_ while active, as breeding buntings may increasingly have to rely on behavioural trade-offs under a warming Arctic to avoid high body temperatures (O’Connor et al., 2022).

## Materials and Methods

### Body and air temperature measurements in calm birds

Buntings were captured on farmlands close to Rimouski, Québec, Canada, and maintained in an outdoor aviary at the Université du Québec à Rimouski as described by Le Pogam et al., (2020). We implanted buntings with a temperature-sensitive, passively integrated transponder (PIT) tag (Biotherm 13, Biomark, Boise USA) either subcutaneously (n = 24) or intraperitoneally (n = 19). Within the subcutaneous group, 6 birds had PIT tags implanted under the left wing on their flank (Nord et al., 2016), while 18 birds had tags implanted on their neck between the scapulae (Oswald et al., 2018). We measured calm (i.e., normal activity) body temperature patterns in a captive population of 43 snow buntings over 3 separate periods: 4 June –12 December 2019, 28 May – 15 November 2020, and 4 March – 16 September 2021. There was no evidence for a difference in temperature between flank and neck locations (*P =* 0.63; mean flank T_b_ = 41.0 ± 1.1 °C and mean neck T_b_ = 41.2 ± 0.7 °C). Consequently, we combined these two locations into one subcutaneous group. Body temperature was recorded every time birds approached the feeder by placing a racket antenna underneath the food source. The racket antenna was attached to a data logger (HPR Plus, Biomark, Boise USA) which scanned for body temperature at 10 sec intervals. Air temperature was recorded at 30 min intervals using iButtons (DS1922L, Maxim Integrated, San Jose USA) suspended 2 m off the ground in the shade.

### Body and air temperature measurements in active birds

We conducted 36 heat stress experiments on 32 actively flying snow buntings from May through December of 2019 and 2020. On the morning of experiments, we measured the body mass of buntings using a digital scale. We timed experiments to coincide with the hottest part of the day, with an average start time of 15:04 ± 0.4 hrs (range = 14:02 – 16:03 hrs). When entering the aviary, buntings begin to fly naturally to avoid capture and therefore experiments started upon entering the aviary (i.e., minute = 0). We captured buntings in flight at various times following minute 0 (range = 32 sec – 33 min) with a butterfly net. Upon capture, we immediately measured the body temperature with the racket antenna and data logger (HPR Plus, Biomark, Boise USA) and subsequently released them into an adjoining aviary. The average duration of experiments was 23 ± 4 min (range = 19 – 33 mins). If buntings had a body temperature ≥ 45°C we immediately submerged their bodies in cold water prior to releasing them to aid in evaporative cooling. Heat stress experiments ended after the last bunting was released and we left the aviary. Air temperature was measured using the same iButtons as for calm birds.

### Data analyses

We performed all analyses in R version 4.1.0 (R Core Team 2022) and reported values are mean ± standard deviation (SD). Mean values represent the combined average across individual bird averages. For example, mean subcutaneous body temperature was calculated by taking a mean body temperature for each bird with a subcutaneous implant and then calculating the combined average across them. We only included body and air temperature values measured between sunrise and sunset for each day (i.e., diurnal body temperatures). We calculated sunrise and sunset times within the *suncalc* package (Thieurmel and Elmarhraoui, 2022). Additionally, we removed all body temperature values recorded during periods when food was replaced and the 10-minute period after leaving the aviary. To place air and body temperature on similar time scales and facilitate comparisons between body and air temperature in calm snow buntings, we rounded each variable to the nearest half-hour. For example, if body or air temperature was measured at 10:44 it was rounded down to 10:30, whereas a recording at 10:45 was rounded up to 11:00. We then averaged body temperature to the nearest 1°C air temperature. For example, mean body temperature at an air temperature of 20 °C included all body temperature values measured between air temperatures of 19.5 and 20.4 °C. We calculated variation in bunting body temperature around their modal body temperature using Boyles et al., (2011a), heterothermy index (HI):

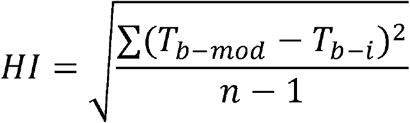

where T_b-mod_ is the modal body temperature in calm birds during daylight hours, T_b-i_, is the body temperature measured at time i, and n is the number of times body temperature was sampled.

Given that modal body temperature represents the most frequently occurring body temperature, we follow Boyles et al., (2011a) and assume this value to represent the optimal body temperature for performance in buntings.

We performed simple linear regression within the *stats* package (R Core Team 2022) when comparing mean body temperature metrics (i.e., mean body temperature, mean modal body temperature, mean maximum body temperature, and mean HI) among subcutaneous and intraperitoneally implanted buntings. When reporting *P*-values, we follow the guidelines of Muff et al., (2022) and present our results in terms of strength of evidence rather than terms of significance. Additionally, we report parameter estimates from all models along with *P*-values.

To determine variation in body temperature patterns among active snow buntings, we used linear mixed-effect models within the *lme4* package (Bates et al., 2015). Bird identity was included as a random factor due to repeated sampling from the same individuals. We first built a global model with the following fixed input variables: air temperature (T_a_), body mass (M_b_), intraperitoneal *vs*. subcutaneous tag location (IP/SQ) and Time (i.e., the time T_b_ was measured in minutes relative to entering the aviary). We also included the two-way interactions between T_a_:M_b_, T_a_:IP/SQ, T_a_:Time, and M_b_:Time, as well as the 3-way interaction between T_a_:M_b_:Time. Prior to fitting the model, we mean-centered and standardized the continuous input variables by dividing by 2 standard deviations (Gelman, 2008) using the standardize function within the *arm* package (Gelman and Su, 2022). We performed model selection on the global model using the dredge function within the *MuMIn* package (Bartoń, 2022) and selected the 95% confidence set of the best-ranked regression models (Symonds and Moussalli, 2011). We then averaged the model parameter estimates from the 95% confidence set. We derived slopes from the model averaged parameter estimates by using the ggpredict function within the *ggeffects* package (Lüdecke, 2018) to calculate estimated marginal means for the response variable. During the 2020 sampling period, we also aimed to quantify the percentage of birds panting relative to air temperature by noting the presence or absence of panting in buntings upon release into the adjoining aviary. Lastly, we calculated the mean hyperthermic scope (T_b_ – T_b-mod_) of indiviual active buntings by subtracting an individual’s measured T_b_ at time of capture from their modal T_b_. All figures were produced using ggplot2 (Wickham, 2016).

## Results

### Body temperature patterns in calm birds

There was substantial variation in mean body temperature across buntings, ranging from 39.4 to 42.6 °C (Figure 1). Body temperatures for both intraperitoneal and subcutaneous implanted birds showed increases above modal body temperature at lower air temperatures (Figure 2). On 19 occasions, 13 different birds (9 intraperitoneal and 4 subcutaneous) had body temperatures ≥ 45°C (Table 1). We found only weak evidence for differences in body temperature among tag location (Table 1).

**Table 1.** Mean body temperature (T_b_), modal body temperature (T_b-mod_), maximum body temperature (T_b-max_) and heterothermy index (HI) across calm, non-stressed snow buntings (*Plectrophenax nivalis*) with intraperitoneal or subcutaneous implants. All values represent diurnal body temperatures recorded between sunrise and sunset. Values in parentheses represent the range of values for individual birds.

| Variable | Intraperitoneal | Subcutaneous | Estimate ± SE | P-value |
| --- | --- | --- | --- | --- |
| Mean $T_b$ | 41.6 ± 0.7 °C<br>(40.4 – 42.6 °C) | 41.2 ± 0.8 °C<br>(39.4 – 42.4 °C) | 0.452 ± 0.229 | 0.055 |
| Mean $T_{b-Mod}$ | 41.6 ± 0.8 °C<br>(39.9 – 42.6 °C) | 41.3 ± 1.1 °C<br>(37.7 – 42.5 °C) | 0.329 ± 0.294 | 0.270 |
| Mean $T_{b-max}$ | 42.2 ± 0.3 °C<br>(39.1 – 51.9 °C) | 41.8 ± 0.3 °C<br>(37.3 – 51.6 °C) | 0.336 ± 0.192 | 0.087 |
| Mean HI | 0.57 ± 0.26°C<br>(0.33 – 1.44°C) | 0.60 ± 0.39°C<br>(0.31 – 1.97°C) | -0.035 ± 0.104 | 0.737 |

**Figure 1.**
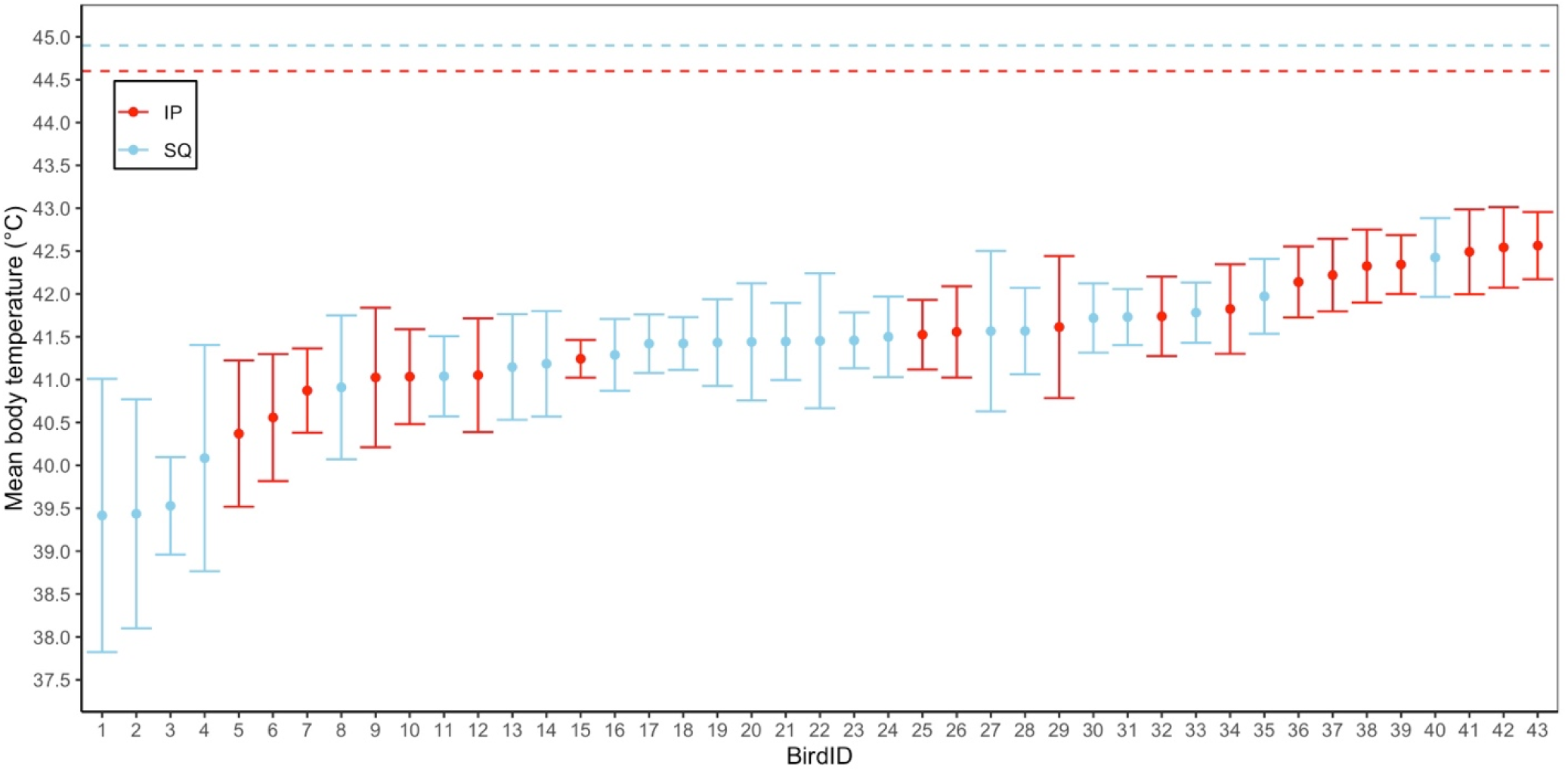
Individual mean body temperatures for calm snow buntings (*Plectrophenax nivalis*) implanted intraperitoneally (IP) or subcutaneously (SQ) with temperature sensitive transponder tags. Error bars represent standard deviation. The horizontal dashed lines are the respective mean maximum body temperatures for intraperitoneal and subcutaneous birds when actively flying. All values represent data recorded between sunrise and sunset.

**Figure 2.**
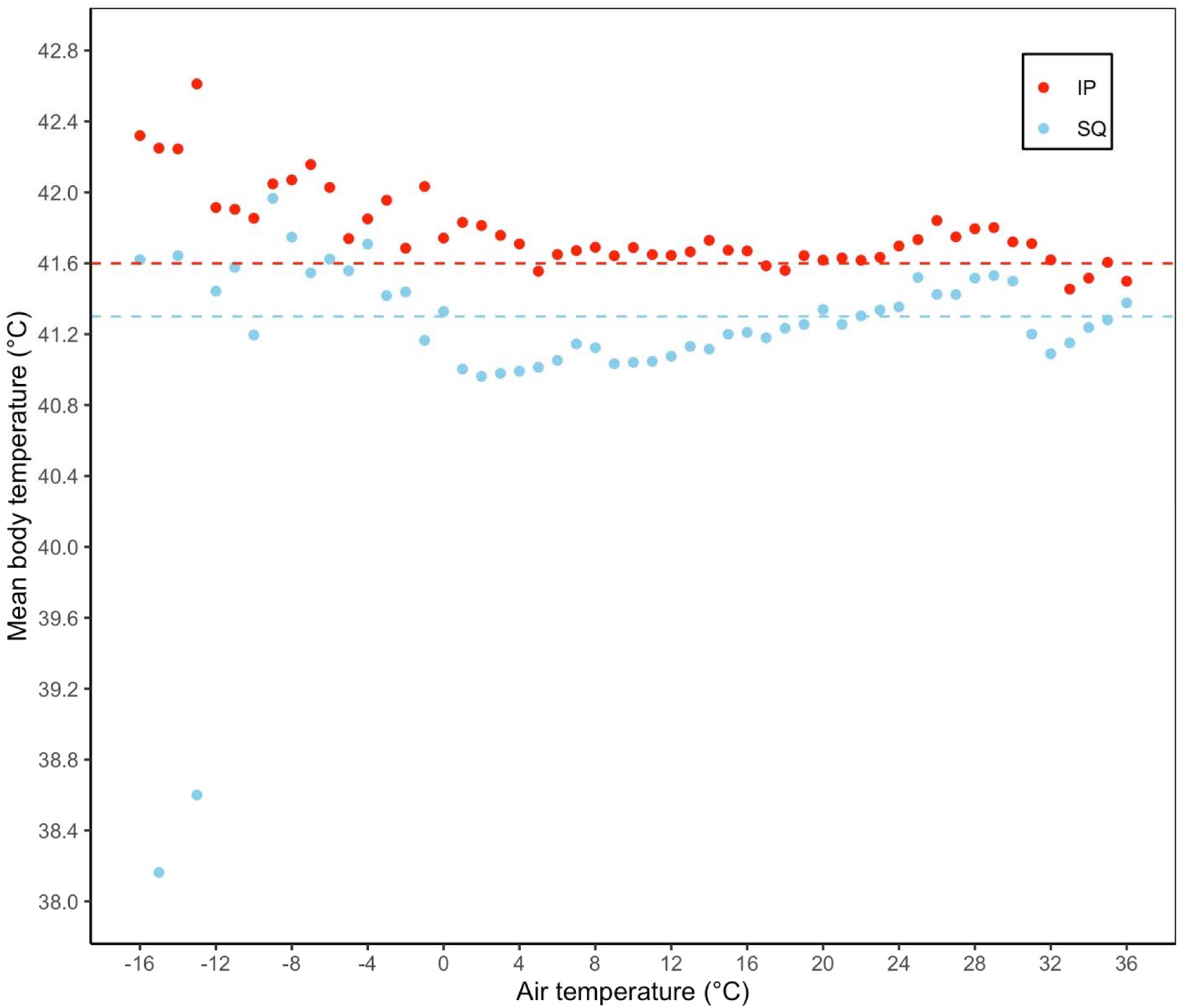
Mean body temperature as a function of air temperature in a captive population of calm snow buntings (*Plectrophenax nivalis*). Body temperature was averaged at 1°C air temperature intervals. For example, mean body temperature at an air temperature of 8°C represents the mean body temperature when air temperatures were between 7.6 and 8.4 °C. Horizontal lines represent the respective average modal body temperatures for intraperitoneal and subcutaneous birds. Error bars were removed for clarity. All values represent data recorded between sunrise and sunset.

### Body temperature patterns in active birds

Active buntings showed a clear and immediate increase in body temperature during experimental trials relative to pre-experiment values (Figure 3a, Table 2). The top candidate model explaining variation in active body temperature among buntings had strong support, being 10x more likely relative to the next best model (Akaike weight of 80.33% *vs*. 8.3%; Table 3). Air temperature, body mass and their interaction were the most important predictive variables given their large effect sizes, large summed Akaike weights, and small *P-*values (Table 4). Body temperature increased linearly with air temperature and on 30 separate experiments, 11 birds (4 intraperitoneal and 7 subcutaneous) had body temperatures ≥ 45°C (Figure 3b). Additionally, heavier birds exhibited a faster increase in T_b_ with T_a_ (i.e., steeper slope). At air temperatures of 9 °C, buntings were already exhibiting body temperatures of 45°C or above (Figure 3b and Figure 4a). At air temperatures from 3.1 °C to 11.1 °C, 27% - 69% of the active birds were panting, while over 50% of the captive bunting population were always panting once air temperatures exceeded 18.6 °C (Figure 4b). Finally, the extent of hyperthermia in buntings was strongly dependent on an individual’s modal body temperature (Estimate = -0.49 ± 0.11 °C, *P* = 0.0001; Figure 5), with mean hyperthermic scope being greater in individuals with lower modal body temperatures.

**Table 2.** Mean body temperature (T_b_), maximum T_b_ (T_b-max_) and the difference between T_b_ and modal body temperature (T_b_ –T_b-mod_) across snow buntings with intraperitoneal or subcutaneous implants approximately 30 min before experimental trials (i.e., calm birds) and during experiments (i.e., actively flying birds).

| Variable | Pre-experiment |  | Experiment |  |
| --- | --- | --- | --- | --- |
|  | Intraperitoneal | Subcutaneous | Intraperitoneal | Subcutaneous |
| Mean $T_b$ | $41.8 \pm 0.6$ °C<br>(40.7 – 42.6 °C) | $41.4 \pm 0.6$ °C<br>(39.8 – 42.5 °C) | $43.5 \pm 0.8$ °C<br>(42.3 – 44.6 °C) | $43.4 \pm 0.8$ °C<br>(41.1 – 44.8 °C) |
| Mean $T_{b-max}$ | $42.4 \pm 0.5$ °C<br>(41.7 – 43.2 °C) | $42.3 \pm 0.6$ °C<br>(41.4 – 44.0 °C) | $44.6 \pm 1.1$ °C<br>(43.2 – 46.9 °C) | $44.9 \pm 0.9$ °C<br>(43.1 – 47.0 °C) |
| Mean $T_b - T_{b-mod}$ | $0.1 \pm 0.7$°C<br>(-0.5 – 2.3 °C) | $0.1 \pm 0.7$°C<br>(-0.7 – 2.1 °C) | $1.9 \pm 0.9$ °C<br>(0.4 – 3.7 °C) | $2.1 \pm 0.7$ °C<br>(1.1 – 3.9 °C) |

**Table 3.**
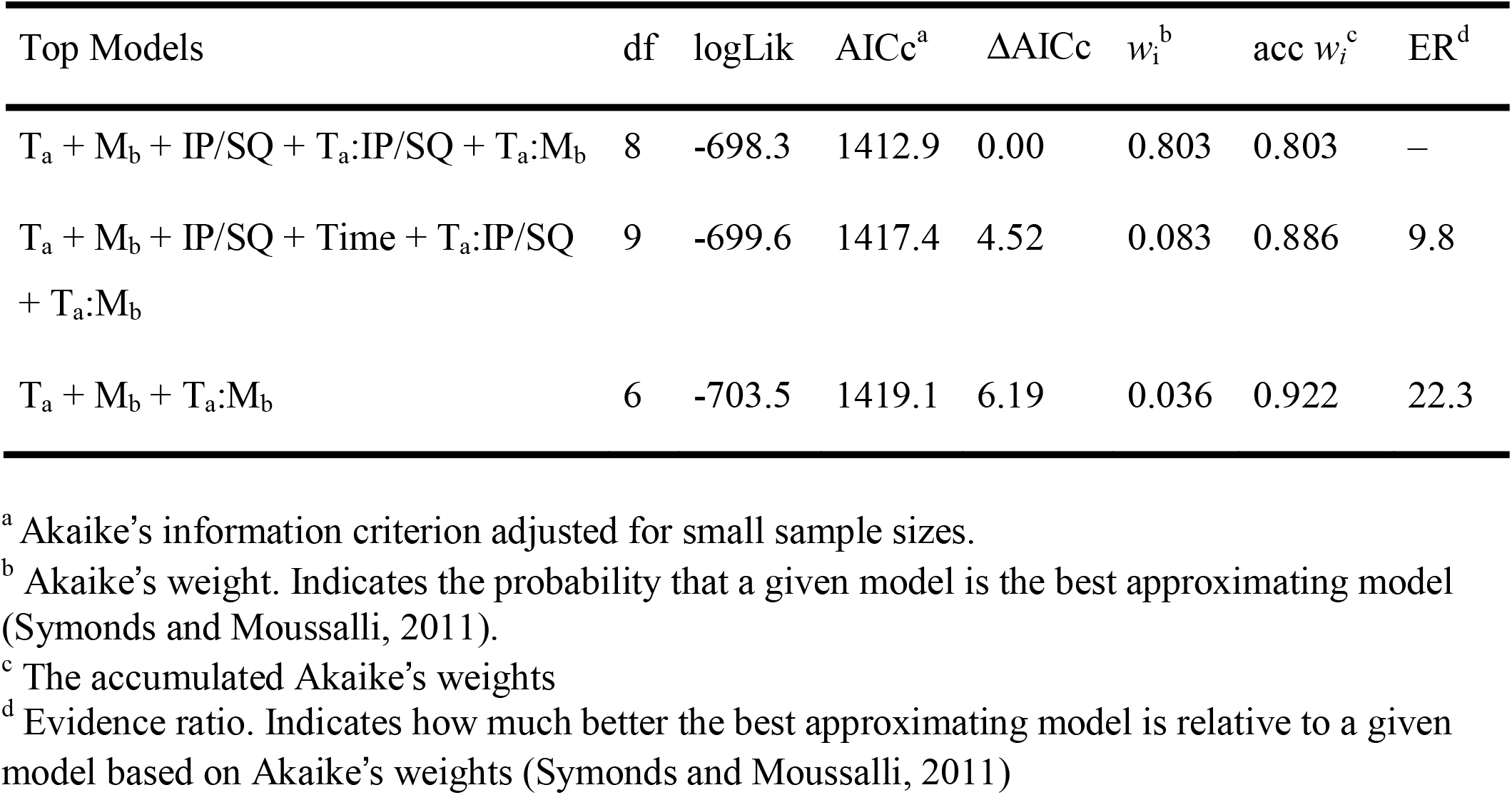
95% confidence set of the best-ranked regression models explaining variation in body temperature among snow buntings during activity experiments. Fixed effects in the top candidate models included air temperature (T_a_), body mass (M_b_), intraperitoneal *vs*. subcutaneous implants (IP/SQ), time of capture (Time), and their interactions.

| Top Models | df | logLik | AICc<sup>a</sup> | $\Delta AICc$ | $w_i^b$ | acc $w_i^c$ | ER<sup>d</sup> |
| --- | --- | --- | --- | --- | --- | --- | --- |
| $T_a + M_b + \text{IP/SQ} + T_a:\text{IP/SQ} + T_a:M_b$ | 8 | -698.3 | 1412.9 | 0.00 | 0.803 | 0.803 | – |
| $T_a + M_b + \text{IP/SQ} + \text{Time} + T_a:\text{IP/SQ} + T_a:M_b$ | 9 | -699.6 | 1417.4 | 4.52 | 0.083 | 0.886 | 9.8 |
| $T_a + M_b + T_a:M_b$ | 6 | -703.5 | 1419.1 | 6.19 | 0.036 | 0.922 | 22.3 |

<sup>a</sup> Akaike's information criterion adjusted for small sample sizes.
<sup>b</sup> Akaike's weight. Indicates the probability that a given model is the best approximating model (Symonds and Moussalli, 2011).
<sup>c</sup> The accumulated Akaike's weights
<sup>d</sup> Evidence ratio. Indicates how much better the best approximating model is relative to a given model based on Akaike's weights (Symonds and Moussalli, 2011)

**Table 4.** Standardized, model-averaged regression estimates from the 95% confidence set of the best-ranked models explaining variation in body temperature among snow buntings during activity experiments.

| Variable | Estimate (±SE) | 95% CI | Summed weight<sup>a</sup> | *P* – value |
| --- | --- | --- | --- | --- |
| Intercept | 43.4 ± 0.1 | 43.1 – 43.7 | NA | <0.0001 |
| T<sub>a</sub> | 0.70 ± 0.06 | 0.58 – 0.82 | 1 | <0.0001 |
| M<sub>b</sub> | -0.90 ± 0.09 | -1.08 – -0.73 | 1 | <0.0001 |
| IP/SQ | 0.39 ± 0.30 | -0.17 – 0.99 | 0.96 | 0.190 |
| Time | 0.006 ± 0.029 | -0.060 – 0.201 | 0.09 | 0.822 |
| T<sub>a</sub>:M<sub>b</sub> | 0.73 ± 0.14 | 0.47 – 1.00 | 1 | <0.0001 |
| T<sub>a</sub>:IP/SQ | -0.42 ± 0.15 | -0.70 – -0.18 | 0.96 | 0.007 |

<sup>a</sup> The summed Akaike’s weights from each model in which that input variable is present. A value of 1 indicates that that variable was in every model within the 95% confidence set of the best-ranked models.

**Figure 3.**
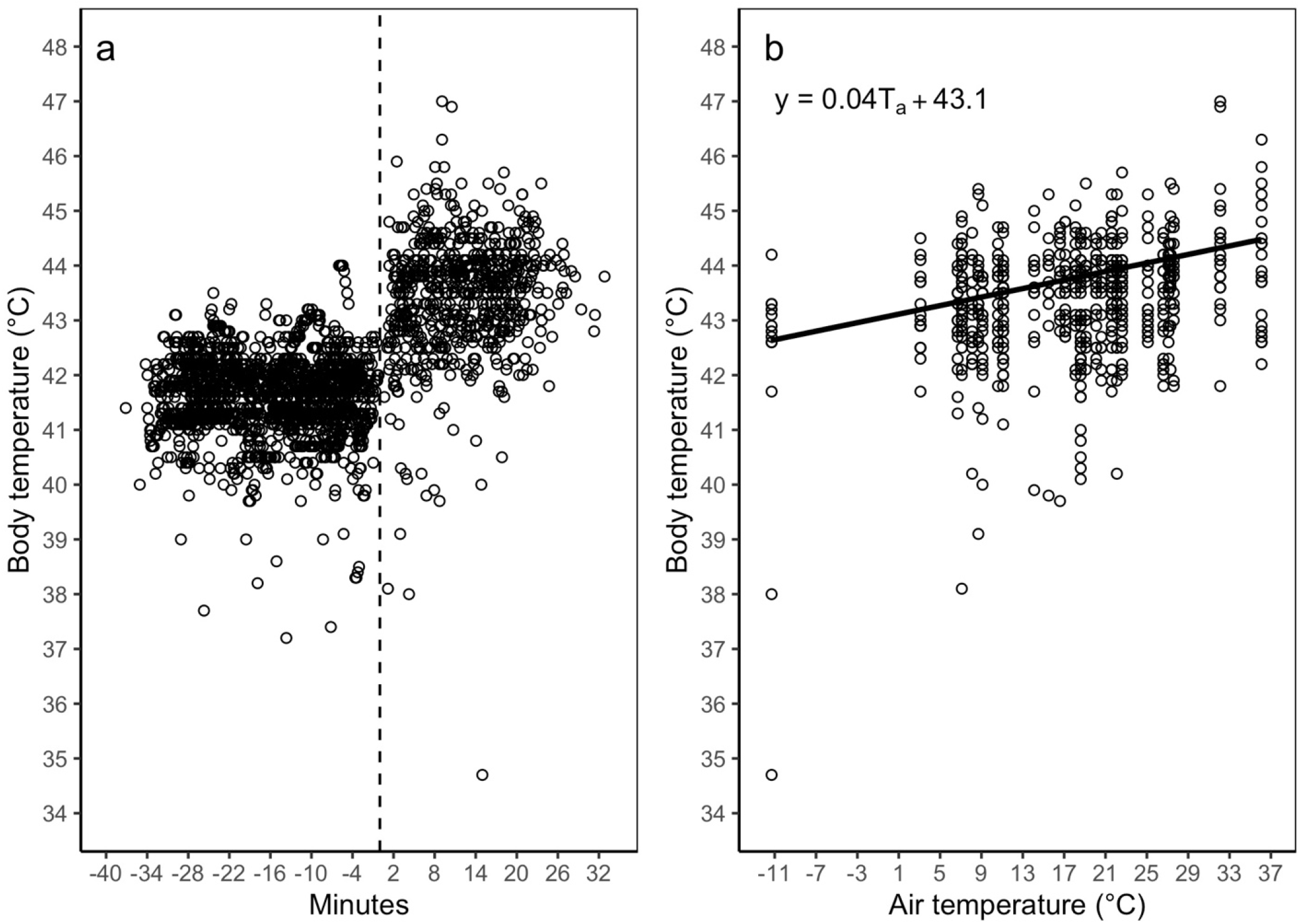
Body temperature patterns in active snow buntings (*Plectrophenax nivalis*) as a function of a) time and b) air temperature. In panel a, negative minutes represent the time span prior to entering the aviary and positive values represent the time span after entering the aviary. The vertical dashed line in panel a) represents the start time for each experiment (i.e., Minutes = 0). In panel b, the regression line represents the marginal effect of air temperature on body temperature from the top model (see Table 1) holding all other input variables constant.

**Figure 4.**
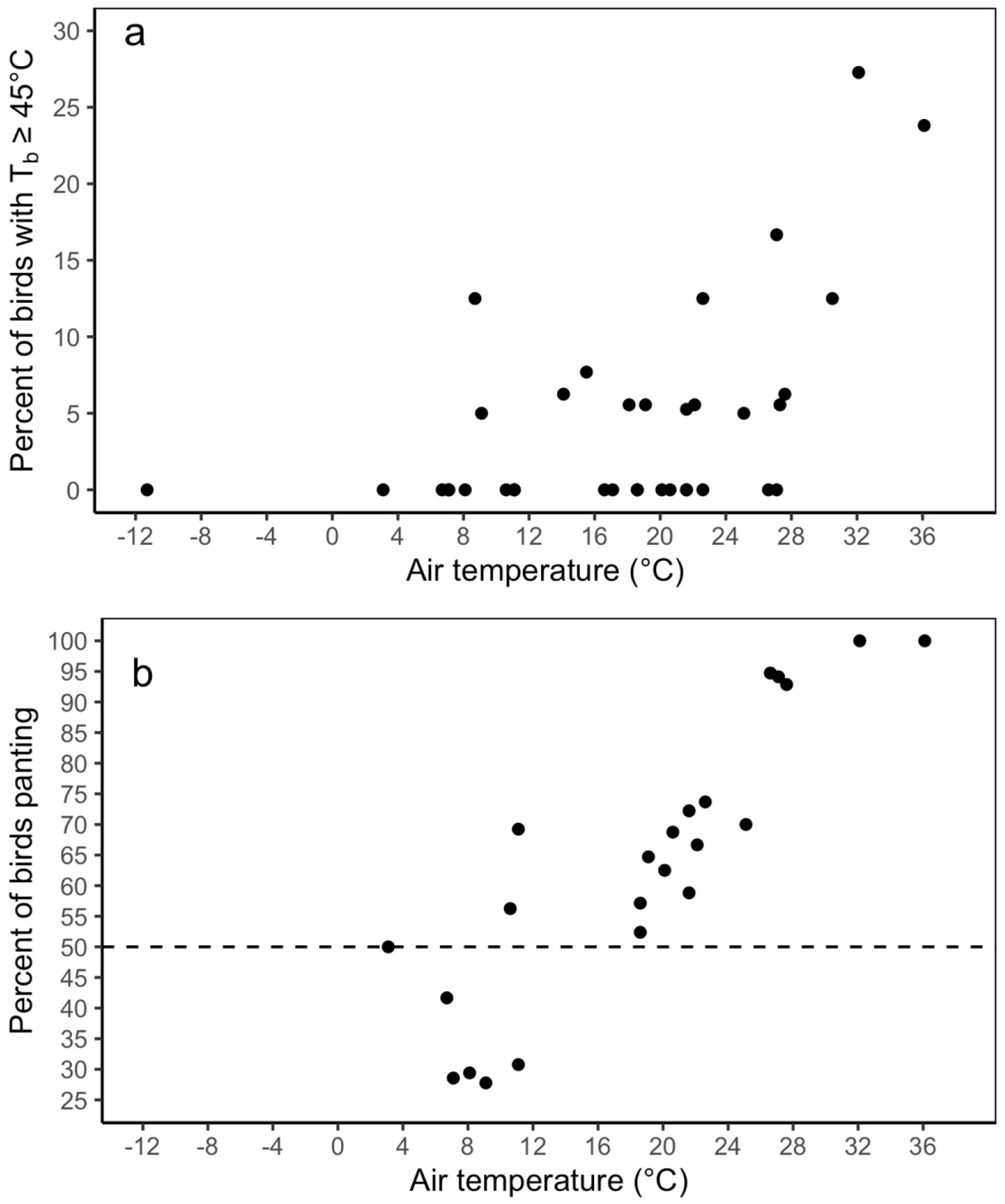
The impacts of air temperature on a) the percent of actively flying snow buntings (*Plectrophenax nivalis*) with a body temperature (T_b_) greater than or equal to 45°C and b) the percent of buntings exhibiting panting behavior. Note the y-axes are on different scales. The fewer data points in panel b) are due to data being collected only during the 2020 study period. The horizontal dashed line in b) represents the threshold above which 50% of the study population was dissipating heat through panting behavior.

**Figure 5.**
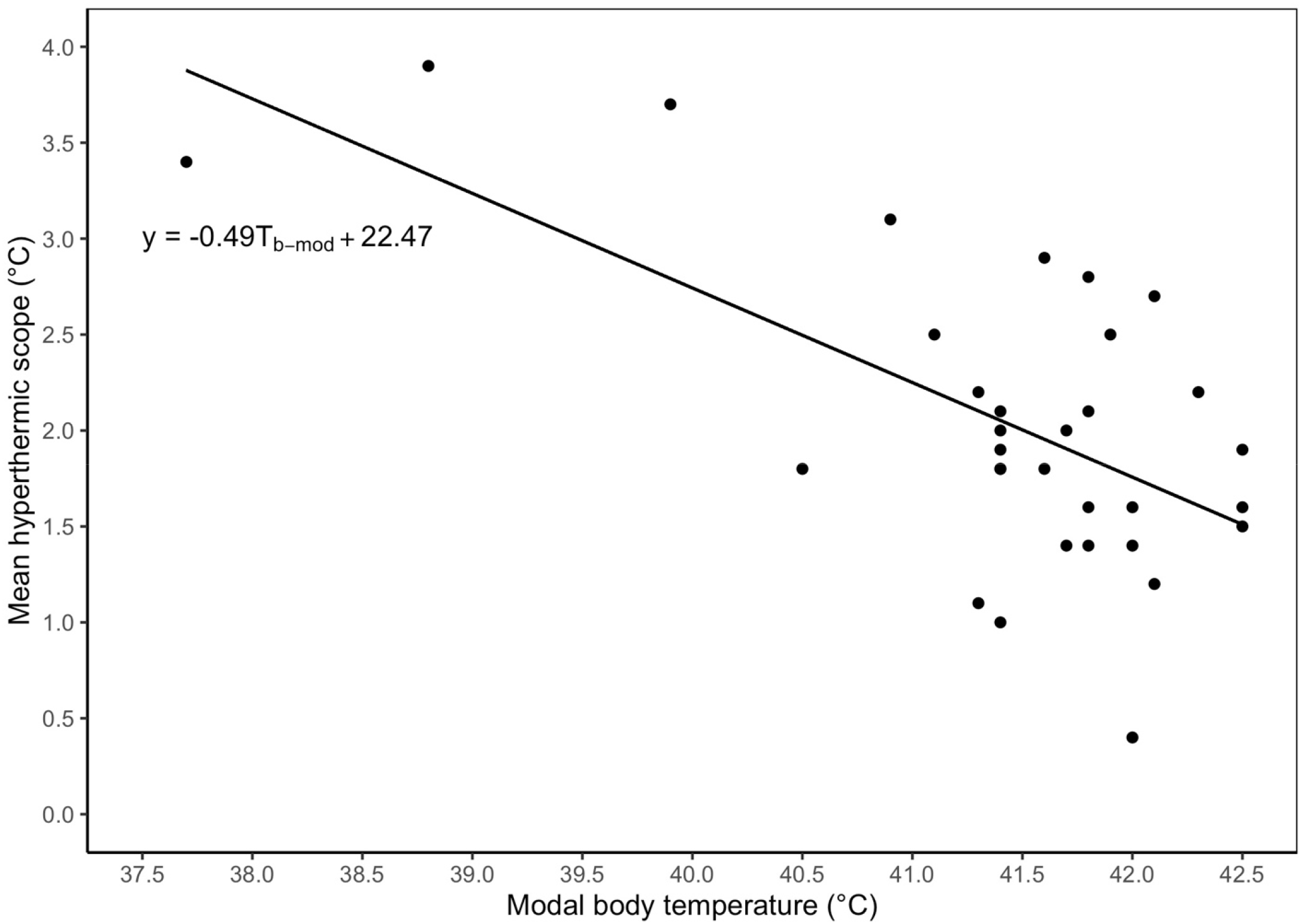
Mean hyperthermic scope (i.e., body temperature – modal body temperature) as a function of modal body temperature (T_b-mod_) among actively flying snow buntings (*Plectrophenax nivalis*).

## Discussion

The interaction between activity and environmental temperature can significantly impact an animal’s body temperature, and concomitantly, thermoregulatory demands, and subsequent behaviour and overall performance (see reviews by Andreasson et al., 2020; Cunningham et al., 2021). Recently, myriad studies originating from hot, arid climates have demonstrated substantial costs on body condition and breeding success arising from behaviourial trade-offs between dissipating heat and the need to perform essential activities (Conradie et al., 2019; Cunningham et al., 2013b; Pattinson and Smit, 2017; Sharpe et al., 2019; van de Ven et al., 2019). Unfortunately, similar data for cold-climate species remains scarce, despite the rapid and unprecendent increases in air temperatures in the Arctic (Rantanen et al., 2022). Here, we sought to help fill this data gap by investigating body temperature increases of actively flying cold-specialist snow buntings in relation to their calm body temperature across a range of environmental temperatures. Specifically, we wanted to determine how maximum body temperature correlated with intense activity under warm air temperatures. At air temperatures of only 9°C, snow buntings already had body temperatures of 45°C or above. Moreover, we observed active snow buntings exhibiting signs of heat stress in the form of panting at very low air temperatures (e.g., < 4°C). These results show that even moderate increases in air temperature can significantly increase a snow bunting’s body temperature when they are active, which in turn increases heat dissipation requirements. Problematically, snow buntings posesses extremely poor evapaorative cooling capacities (O’Connor et al., 2021) and therefore will likely rely heavily on reducing their activity to thermoregulate (O’Connor et al., 2022). As a result, our findings strongly indicate that as the Arctic continues to warm, snow buntings will increasingly rely on behavioural adjustments in activity to avoid lethal body temperatures during energetically expensive stages such as the nestling-provisioning period. The culmination is expected to be substantial costs to nestling condition and even nestling survival, ultimately impacting breeding success.

The nestling-provisioning period for birds is considered one of the most energetically demanding life-history stages in altricial species (Drent and Daan, 1980). However, as air temperatures increase under climate change, sustaining the optimal work levels required for feeding nestlings will become increasingly harder as individuals will likely experience higher body temperatures requiring increased heat issipation behaviours (O’Connor et al., 2022).

Indeed, our data show that buntings regularly experienced higher body temperatures at higher air temepratures, frequently exceeding 45°C, a value approaching lethal levels (Freeman et al., 2020; McKechnie and Wolf, 2019). Similarly, (Nilsson and Nord, 2018) found that nestling-provisioning marsh tits (*Poecile palustris*) originating from a cool-temperate climate had higher body temperatures on days with higher ambient temperatures, with body temperatures also exceeding 45°C on multiple occassions. The cause for increased body temperatures among buntings in response to increasing air temperature is likely two-fold. Firstly, snow buntings are known cold-specialists having evolved physiological traits for toleraing extreme cold (Le Pogam et al., 2020), possibly at the detriment for heat tolerance. Secondly, as air temperatures increase, the thermal gradient between body and air temperature becomes increasingly narrow, thereby restricting the flow of heat through non-evaporative pathways. Given that snow buntings are known to be extremely limited in their capacity to evaporatively dissipate heat (O’Connor et al., 2021), it becomes increasingly harder for them to dissipate heat at higher ambient temperatures. Moreover, recent work has shown that buntings maintain cold hardniness into both northward migration and even after arrival on the breeding grounds (Le Pogam et al., 2021a; Le Pogam et al., 2021b). Collectively, these results underscore the likelihood that as the Arctic continues to warm, nestling-provisioning buntings performing at high sustained metabolic rates will increasingly experience periods where they are unable to physiologically limit their body temperature from approaching lethal levels, requiring buntings to behaviourally allocate more time towards dissipating heat. As has recently been demonstrated for hot, arid bird species (Conradie et al., 2019; Cunningham et al., 2021; Smit et al., 2016), behavioural adjustments are likely to result in a cost to essential parental activities (e.g., nestling feeding) and therefore fitness (O’Connor et al., 2022).

The ability for ecologists to predict temperature theresholds above which species will begin to experience fitness costs associated with climate change remains a significant challenge (Angilletta and Sears, 2011; Cunningham et al., 2013a; Schwenk et al., 2009). Although valuable, mechanistic models used to calculate mass and heat transfer rates among species are complex, requiring numerous physiological and morphological parameters. Recently, Smit et al., (2016) proposed a less computationally intense heat dissipation behaviour index for assessing the vulnerability of species to high environmental temperatures. Specifically, this index uses presence/absence data to calcaulate when heat dissipation behaviours (e.g., panting or wing spreading) occur in at least 50% of instances. Using this index, Smit et al., (2016) found that among 33 Kalahari desert bird species, the median air temperature at which panting occurred in 50% of observations ranged from approximately 30 °C to 46 °C. In stark contrast, we found that by air temperatures ranging from 12 °C to 18 °C, 50% or more of our snow buntings were exhibiting panting behavior. The lower end of this air temperature range aligns remarkably well with O’Connor et al.’s 2022) recent prediciton that at an air temperature of only 11.7 °C buntings in the Arctic will begin to experience behavioral trade-offs stemming from increased thermoregulatory demands. However, O’Connor et al.’s (2022) prediction was based on a sustainable metabolic rate maintained at 4-times basal metabolism during nestling provisioning, and snow buntings in the current study were likely exerting higher metabolic rates. Regardless, these findings ultimately show that buntings must increase their thermoregulatory behavior in response to increases in T_b_ at moderately low air temperatures, suggesting they will certainly experience increased thermoregulatory demands on their Arctic breeding grounds in the future.

The regulation of body temperature can significantly influence the transfer of heat and thus affect an animals thermoregulatory demands (Angilletta and Cooper, 2010; Boyles et al., 2011b; Scholander et al., 1950; Tattersall et al., 2012). Indeed, most birds have been shown to facultatively increase their body temperature when subjected to high air temperatures, an adaptive trait often attributed to enhancing water economy (Weathers 1981, Tieleman and Williams 1999, Gerson et al. 2019). Consequently, although seemingly counterintuitive, the regulation of a lower T_b_ may be advantageous in the presence of high air temperautres as ones baseline T_b_ is further away from lethal levels, therefore lessening the risk of overheating when becoming hyperthermic (McNab and Morrison, 1963). Overwhelmingly, investigators have studied adaptive thermoregulation among populations in response to climatic variables (e.g., arid versus mesic), both at the inter-and intra-specific level (Noakes and McKechnie, 2019; Smit et al., 2013; Tieleman, 2007). However, to our knowledge no studies have reported intra-specific body temperature variation within the same population. Here, we found that mean body temperature and modal body temperature among snow buntings varied by 2.2°C and 2.7°C, respectively. Moreover, we found strong evidence that individuals with higher modal T_b_ at rest showed smaller increases in T_b_ when active, suggesting that birds with a higher resting T_b_ (i.e., when calm) may have a reduced scope for further T_b_ increases when performing active portions of their life history. While further work is required to determine whether these differences in scope are fixed at the individual level, variation in a set-point body temperature within the same population could allow certain individuals to prolong activity at high air temperatures given the larger spare capacity for hyperthermia. Consequently, buntings with a lower set-point body temperature could have a selective advantage under a rapidly warming Arctic as they would be less at risk of overheating when working at peak levels. Of important note, to our surprise, snow buntings maintained higher body temperatures at colder environmental temperatures. This pattern is contrary to typical patterns, wherein body temperature typically remains stable across a range of environmental temperatures below the thermoneutral zone but begins to increase with increasing environmental temperatures within and above the thermoneutral zone (Tieleman and Williams, 1999; Weathers, 1981). Currently, the reason for our findings is not yet clear and the underlying driver(s) of this pattern requires more examination in this and possibly other winter-acclimated passerine species.

